# Measures of Possible Allostatic Load in Comorbid Cocaine and Alcohol Use Disorder: Brain White Matter Integrity, Telomere Length, and Anti-Saccade Performance

**DOI:** 10.1101/347138

**Authors:** Jonika Tannous, Benson Mwangi, Khader M. Hasan, Ponnada A. Narayana, Joel L. Steinberg, Consuelo Walss-Bass, F. Gerard Moeller, Joy M. Schmitz, Scott D. Lane

## Abstract

Chronic cocaine and alcohol use impart significant stress on biological and cognitive systems, resulting in changes consistent with an allostatic load model of neurocognitive impairment. The present study measured potential markers of allostatic load in individuals with comorbid cocaine/alcohol use disorders (CUD/AUD) and control subjects. Measures of brain white matter (WM) integrity, telomere length, and impulsivity/attentional bias were obtained. WM integrity (CUD/AUD only) was indexed by diffusion tensor imaging metrics, including radial diffusivity (RD) and fractional anisotropy (FA). Telomere length was indexed by T/S ratio. Impulsivity and attentional bias to drug cues were measured via eye-tracking, and were also modeled using the Hierarchical Diffusion Drift Model (HDDM). Average whole-brain RD and FA were associated with years of cocaine use (R^2^ = 0.56 and 0.51, both p < .005) but not years of alcohol use. CUD/AUD subjects showed more anti-saccade errors (p < .01), greater attentional bias scores (p < .001), and higher HDDM drift rates on cocaine-cue trials (Bayesian probability CUD/AUD > control = p > 0.99). Telomere length was shorter in CUD/AUD, but the difference was not statistically significant. Within the CUD/AUD group, exploratory regression using an elastic-net model determined that more years of cocaine use, older age, larger HDDM drift rate differences and shorter telomere length were all predictive of white matter integrity as measured by RD (model R^2^ = 0.79). Collectively, the results provide modest support linking CUD/AUD to putative markers of allostatic load.

## Introduction

Allostasis implies a shift in homeostatic systems in response to acute or chronic stressors (e.g., allostatic load), described by McEwen as “the price the body pays for being forced to adapt to adverse psychosocial or physical situations” [1]. Exposure to extreme or pervasive stressors can result in pathophysiological change. Examination of allostatic load has commonly focused on changes in the regulation of stress hormones and the sequelae of such changes [2,3], but the concept can apply broadly to effects on many homeostatic systems. Nearly two decades ago, Koob and colleagues proposed allostatic models of chronic cocaine and alcohol intake based on preclinical work [4–6], but only modest attention has focused on the allostasic framework in studying human substance use disorders (SUD). Here, we focus on the effects of chronic SUD in adults with co-morbid cocaine and alcohol use disorders [7–9].

Cocaine use disorder (CUD) with co-occurring abuse of other substances, e.g., marijuana and alcohol use disorder (AUD), is the norm rather than the exception [10]. Individuals who use only cocaine are of putative scientific importance with regard to isolating individual drug effects. However, such investigations are more precisely understood using preclinical models that can isolate dose-response relationships. In reality, because cocaine-only users are rare, they may represent a phenotype of lesser clinical or applied interest. From the standpoint of measuring indicators of CUD that translate to real world effects, alterations in biological and neurocognitive function are probably best understood as the result of the synergistic impact of CUD plus abuse of other substances.

The present study focused on brain white matter integrity, telomere length, and eye movement indices of impulse/attentional control. While not a comprehensive set of biological or cognitive markers of CUD/AUD-related impairment, these domains are indicators of health and cognitive functioning. Importantly, varying degrees of evidence exist to suggest that each domain is adversely modified by chronic cocaine and alcohol use [11–18]. Each marker – white matter integrity, telomere length, and anti-saccade performance – is broadly associated with neurological and psychiatric disease processes, independent of substance use disorders [19–30], and thus collectively provides an indicator of possible allostatic load extrapolated from a broader evidence base applied to CUD/AUD. We expected to observe evidence of impairment for each marker related to CUD/AUD and the cumulative effects of abusing these substances.

## Methods

### Subjects

Participants for this project were recruited from the Greater Houston Metropolitan Area using local newspaper and radio advertisements. The data reported here constitute part of a larger clinical trial examining the effects 12-weeks of treatment with pioglitazone in participants with a primary cocaine use disorder (CUD) and a secondary alcohol use disorder (AUD), described in [31] (NCT02774343). For the present report, data were obtained from measures taken at baseline (Day 0), prior to initiation of the clinical trial, and included 22 CUD/AUD subjects who provided complete neuroimaging, eye tracking, and telomere data; two additional CUD/AUD subjects provided eye tracking and telomere data without DTI. In addition, data from two independent samples of 35 (eye tracking) and 25 (telomere) healthy control subjects were obtained for purposes of comparison with the CUD/AUD subjects. DTI was not obtained from control participants, who were not part of the clinical trial, as neuroimaging of control subjects was beyond both the scope and budget of the project. Additionally, numerous studies have previously characterized white matter integrity in CUD and AUD [11–13,32–35]. The study was carried out in accordance with the recommendations of the Belmont Report and the approval of the UTHSC-Houston IRB. All subjects gave written informed consent obtained in person in accordance with the Declaration of Helsinki. Subject characteristics are provided in Table 1.

**Table 1.** Demographic characteristics for CUD/AUD subjects (N=22 completed DTI) and control groups for the anti-saccade (eye-tracking) and telomere length (telomere) data. Values represent mean (SD) or Number (#, %).

| GROUP | CUD/AUD | control (eye-tracking) | control (telomere) |
| --- | --- | --- | --- |
| N | 24 | 35 | 25 |
| Age | 46.96 (7.66) | 42.49 (10.17) | 43.76 (6.62) |
| Sex: # Male (%) | 18 (75.00) | 17 (48.57) | 17 (68.00) |
| Education | 12.69 (1.52) | 14.06 (2.39) | 14.00 (2.55) |
| Race: # (%) |  |  |  |
| African Am | 14 (58.33) | 25 (71.43) | 18 (72.00) |
| Caucasian | 6 (25.00) | 5 (14.29) | 2 ( 8.00) |
| Hispanic | 4 (16.67) | 5 (14.29) | 5 (20.00) |
| Shipley | 87.67 (13.10) | 94.63 (14.15) | 99.60 (10.87) |
| Cocaine (yrs) | 17.96 (8.34) | -- | -- |
| Alcohol (yrs) | 21.62 (12.27) | -- | -- |

### Measures

#### Demographic and Clinical

The following measures were obtained for all subjects: age, sex, education, race, cognitive aptitude (Shipley II, [36]), and mental health functioning via Structured Clinical Interview for DSM-IV (SCID-IV, [37]). For CUD/AUD subjects, lifetime and recent substance use were determined via the Addiction Severity Index [38], and the Kreek-McHugh-Schluger-Kellogg scale (KMSK, [39]).

#### Diffusion Tensor Imaging (DTI)

DTI is a magnetic resonance imaging technique that is used to map brain white matter (WM) fibers by quantifying the tissue diffusion properties of water. DTI-derived metrics examined here included axial, radial, and mean diffusivity (AD, RD, and MD, respectively), as well as fractional anisotropy (FA). These metrics have been used to infer CNS white matter pathology in a number of neurological and psychiatric diseases [40-44]. DTI methods, scanning parameters, and data processing details are provided in Supplement S1 Text. In summary, for each subject (N = 22) whole brain voxel-wise FA, MD, RD, and AD maps were generated with toolboxes provided by FSL [45] using the Tract-Based Spatial Statistics (TBSS) method [46,47]. DTI-derived maps were transformed into MNI152 standard space by nonlinearly registering each map to a standard template provided with FSL using a non-linear registration method (FNIRT, [47]). Each subject’s FA map was projected onto the corresponding FA skeleton, allowing for voxelwise analysis across subjects using a permutation-based non-parametric statistical method (RANDOMISE, [48]) with 5000 permutations. T1-based analyses with SPM and Freesurfer examining grey matter volume, white matter volume, and cortical thickness and their relationships to cocaine use did not reveal any significant patterns. Additionally, all scans were reviewed by a radiologist to ensure that there were no gross anatomical abnormalities. Consequently, the present DTI-based results may represent a unique predictor of potential changes not be captured with volumetrics or brain abnormalities.

#### Anti-Saccade Task

The details of the eye-tracking task are provided in Dias et al [15]. Briefly, participants were tested using an infrared binocular eye-tracker to measure performance on blocks of pro-saccade and anti-saccade trials (36 trials per block). Following training to minimize blinking and optimize task understanding, four counterbalanced blocks were administered (2 pro-saccade, 2 anti-saccade, 144 trials total). Stimuli included cocaine-related images, neutral images matched as closely as possible to the cocaine images (e.g., color, background, complexity), and size-matched solid gray shapes. The eye-tracking data were analyzed as (1) overall error rates on all anti-saccade trials (global inhibitory control), (2) the ratio of anti-saccade errors on cocaine-stimulus trials to total anti-saccade errors over all trials (cocaine/total: attentional bias towards cocaine cues). Anti-saccade errors provide an index of inhibitory control circuitry, both with regard to neural pathways subserving the control of eye movements and to markers of pathology in psychiatric and neurological disease [49,50]. Attentional bias provides an index of asymmetrical attentional control by the salience of specific stimuli (in this case cocaine cues) relative to neutral stimuli, with clinical relevance to SUD [15,51,52].

#### Hierarchical Drift Diffusion Model (HDDM)

Error rates and attentional bias are robust but coarse and descriptive measures of anti-saccade performance. Accordingly, eye-tracking performance was also examined with a conceptually-informed approach from cognitive neuroscience, the HDDM [59]. The HDDM is a drift diffusion / sequential sampling model of decision making that incorporates both accuracy and reaction time, based on a threshold model in which evidence for a decision (here, execution of an anti-saccade) accumulates in the context of imperfect information (noise). The HDDM capitalizes on both the theoretical utility of the drift diffusion framework and the use of Bayesian data analytic methods, which have substantial utility in modeling neural and cognitive phenomenon [53]. The analysis focused on the HDDM parameter *v* (drift rate). Herein, we interpret differences in *v* as evidence of differential salience of the cocaine (vs. non-drug) stimuli, such that for the CUD/AUD group the cocaine stimuli increase decision conflict reflected in both accuracy and reaction time. Details of this model, including conceptual foundations and neural correlates can be found in [54,55]. Additionally, details of the HDDM outcomes, including model output (group and individual subject), and metrics of model convergence and fit are provided in Supplement S2 Data.

#### Telomere Length

Genomic DNA was extracted from leukocytes by standard procedures. DNA concentration was assessed by Nanodrop and telomere length was measured quantitatively by PCR, as previously described [56]. Briefly, primers for the telomere sequence (T) are tel1b: 5’- CGGTTT(GTTTGG)5GTT-3’ and tel2b: 5’-GGCTTG(CCTTAC)5CCT-3’. The single-copy gene human beta-globin was used as the reference gene (S), with the following primers: hbg1 5’-GCTTCTGACACAACTGTGTTCACTAGC-3’ and hbg2 5’ CCAACTTCATCCACGTTCACC-3’. T and S values were quantified relative to a reference DNA sample by the standard curve method. Since the number of S copies are the same in all individuals, relative T/S ratio (the primary dependent variable) reflects relative length differences in telomeric DNA. All PCRs were carried out using the thermal cycling profiles previously described in [56], with DNA samples run in duplicate on separate plates but in the same well positions.

### Data Analytic Strategy

#### DTI: Confirmatory Analyses

We examined the relationship between FA, RD, MD and AD values and years of cocaine use while controlling for alcohol use (and vice versa), and combined years of both substances (adjusting for age and gender in all models). As no control groups were available for DTI, all analyses were within CUD/AUD subjects. Combined years of use was calculated by adding the years of cocaine use to years of alcohol use. Based on clinical interview at screening, including the ASI, SCID, and the Kreek-McHugh-Schluger-Kellogg (KMSK) scale, cocaine and alcohol use were consistent across individual subjects’ lifetime (e.g., no marked periods of abstinence or changes in use patterns). Therefore, given subjects’ age range and history of use (cocaine mean = 5.13, range = 5.5 – 36.0 years; alcohol mean = 6.18, range = 2.75 – 40.0 years), we determined that years of use was the best proxy for cocaine and alcohol exposure with regard to recall accuracy, minimizing error in the measurement of extent of substance exposure. Notably, chronicity has been cited as a critical component in addiction diagnostics [57]. All regression models adjusted for age, education, sex and Shipley score, with the significance threshold set at *p* < 0.005. The analyses resulted in a map highlighted by all significant clusters. To visualize the relationships found, the average FA value across significant clusters was calculated for each subject and then plotted against years of use. The same pipeline was then used for the MD, RD, and AD scalars.

#### Anti-Saccade Task: Confirmatory Analyses

Consistent with Dias et al [15] and with the overall literature on saccade-based performance, there were no differences between CUD and control subjects on pro-saccade trials. Accordingly, data analyses focused on anti-saccade trials. Per Pocock [58], we examined the influence of potential confounding variables (age, education, sex, Shipley score) by testing for (1) between-group differences and (2) correlations with the dependent variables of interest (total anti-saccade errors, attentional bias). Age and education showed group differences and correlations with *p*-values < .10, and were thus treated as covariates in linear regression models that examined differences between CUD/AUD vs. controls on total anti-saccade errors and attentional bias.

#### HDDM: Confirmatory Analyses

The HDDM package in Python [54,59] was used to fit a hierarchical model to compare the probability distributions of differences in *v* (drift rate) between groups and stimulus types. The HDDM package provides estimation of differences in Bayesian posterior probability distributions by computing the proportion of posteriors in which *v* (drift rate) is greater in one condition than another. Differences were compared for each stimulus type (cocaine, neutral, shape) both within and between groups. Model fitting and convergence details are provided in Supplement S2 Figs.

After comparing the Bayesian probability distributions of differences in *v* (drift) between groups and between stimuli, individual *v* (drift) values for each subject were extracted and entered into a linear regression model controlling for age and education, in order to replicate the confirmatory frequentist models used to analyze total anti-saccade errors and attentional bias. To create a single dependent variable for the frequentist analyses, a difference score was created for each subject by subtracting *v* on cocaine-stimulus trials from non-cocaine stimulus (neutral+shape) trials.

#### Telomere Length: Confirmatory Analyses

The T/S ratio was examined in a linear regression model that compared differences between CUD/AUD vs. controls with age and education as covariates in same manner as described above.

#### Hypothesis-Generating Analyses: White Matter, Anti-saccades, and Telomere Length

Because the dataset was limited to 22 subjects who provided complete DTI, eye-tracking and telomere data, we utilized modern regularization techniques with penalized regression. Two primary penalized regression approaches, ridge and lasso, can be linearly combined via elastic-net regression, which overcomes limitations of each type but includes both as special cases [60,61]. Leave-one-out cross-validation (LOOCV) was employed to optimize the tuning parameters of the model, e.g., to determine the alpha (the mixing or penalty parameter, range 0-1) and lambda (regularization or coefficient shrinkage parameter) parameters of the elastic net model that minimize the mean squared error [61,62]. These analyses were conducted using the R glmnet package, based on modeling techniques recommended by the authors [60]. To provide further model interpretability, the R package selectiveInference [63] was integrated with the glmnet model outcomes to provide z-score, p-value, and confidence interval estimates for the obtained model coefficients, as per [62]. Using this approach, we modeled the relationship between DTI radial diffusivity (RD, dependent variable) and the following predictors: telomere length (T/S ratio); anti-saccades (HDDM difference score on *v* for cocaine vs. non-cocaine trials); and total years of cocaine use. To control for relevant covariates, age and years of alcohol use were included in the model. RD was selected because it contained the largest number of significant clusters related to years of cocaine use, and because of its probable connection to compromised myelination [11,32,64]. All variables were standardized by z-scoring to provide interpretability of the model coefficients.

## Results

### Diffusion Tensor Imaging

As shown in Fig 1, FA values decreased as a function of years of cocaine use, controlling for years of alcohol use, age, sex, education and Shipley score, p < .005, R^2^ = 0.56. The significant clusters (in yellow) associated with cocaine use were identified along several major tracts (in green), including the corpus callosum, the right thalamic radiation, the right superior longitudinal fasciculus, and the corona radiata. Notably, analysis of AD values revealed no significant relationship between AD and years of cocaine use, while RD and MD values increased with years of cocaine use, R^2^ = 0.51 and 0.46,respectively. RD clusters were the most widespread and included the internal capsule, corona radiata, optic radiation, tapetum, superior longitudinal fasciculus, posterior thalamic radiation, and all clusters also found in the FA maps. Although there were significant RD clusters in both hemispheres, clusters were larger in the right hemisphere. Fig 1 highlights the significant FA clusters, and provides a scatterplot of the relationship between FA and years of cocaine use, controlling for covariates. Supplement S3 Table provides details for all significant clusters for the FA, RD, and MD maps.

**Fig 1.**
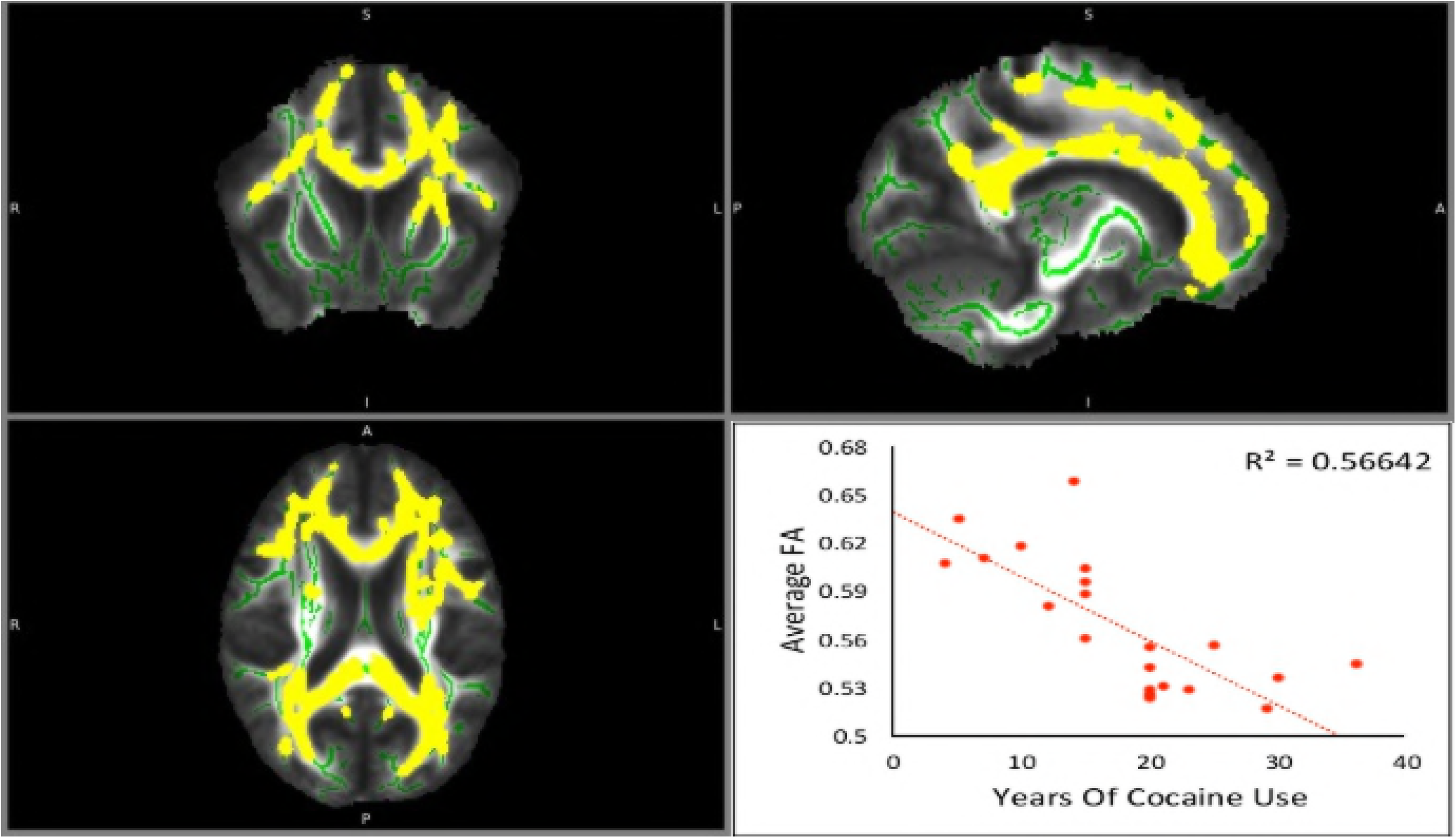
Fractional anisotropy (FA) maps, showing coronal, sagittal, and axial views (respectively) of white matter tracts in 22 CUD/AUD subjects. Major tracts are outlined in green and areas highlighted in yellow represent group-level significant associations with years of cocaine use. The bottom right panel shows a scatter plot of average whole-brain FA value (y-axis) and a function of years of cocaine use (x-axis) for individual subjects (R^2^ = 0.566.)

No significant clusters were found when examining the relationship between any of the DTI scalars with years of alcohol use or years of combined cocaine + alcohol use, controlling for other relevant covariates. With no observed relationships within the AD values, it follows that the significant clusters in the FA map are probably driven by the relationship between RD and years of cocaine use. Consequently, all of the areas significant in the FA and MD maps are also significant in the RD map. The RD map contains the largest number of significant clusters, which were not lessened by input from non-significant AD differences.

### Anti-Saccade Task

Fig 2 summarizes the total anti-saccade error and attentional bias data. For the eye-tracking data linear models, degrees of freedom and R^2^ values are provided along with t-values, p-values, and group means (± SEM) for any significant predictor variables. For total anti-saccade errors, df = 3, 55, R^2^ = 0.24: group t = 2.70, p < .001; age t = 2.31, p < .03. CUD/AUD made more anti-saccade errors than controls, 28.04 (± 2.96) vs. 17.77 (± 2.34). For attentional bias scores, df = 3, 55, R^2^ = 0.29: group t = 4.69, p < .001; age was *ns*. CUD/AUD had higher attentional bias scores than controls, 0.47 (± 0.03) vs. 0.32 (± 0.03).

**Fig 2.**
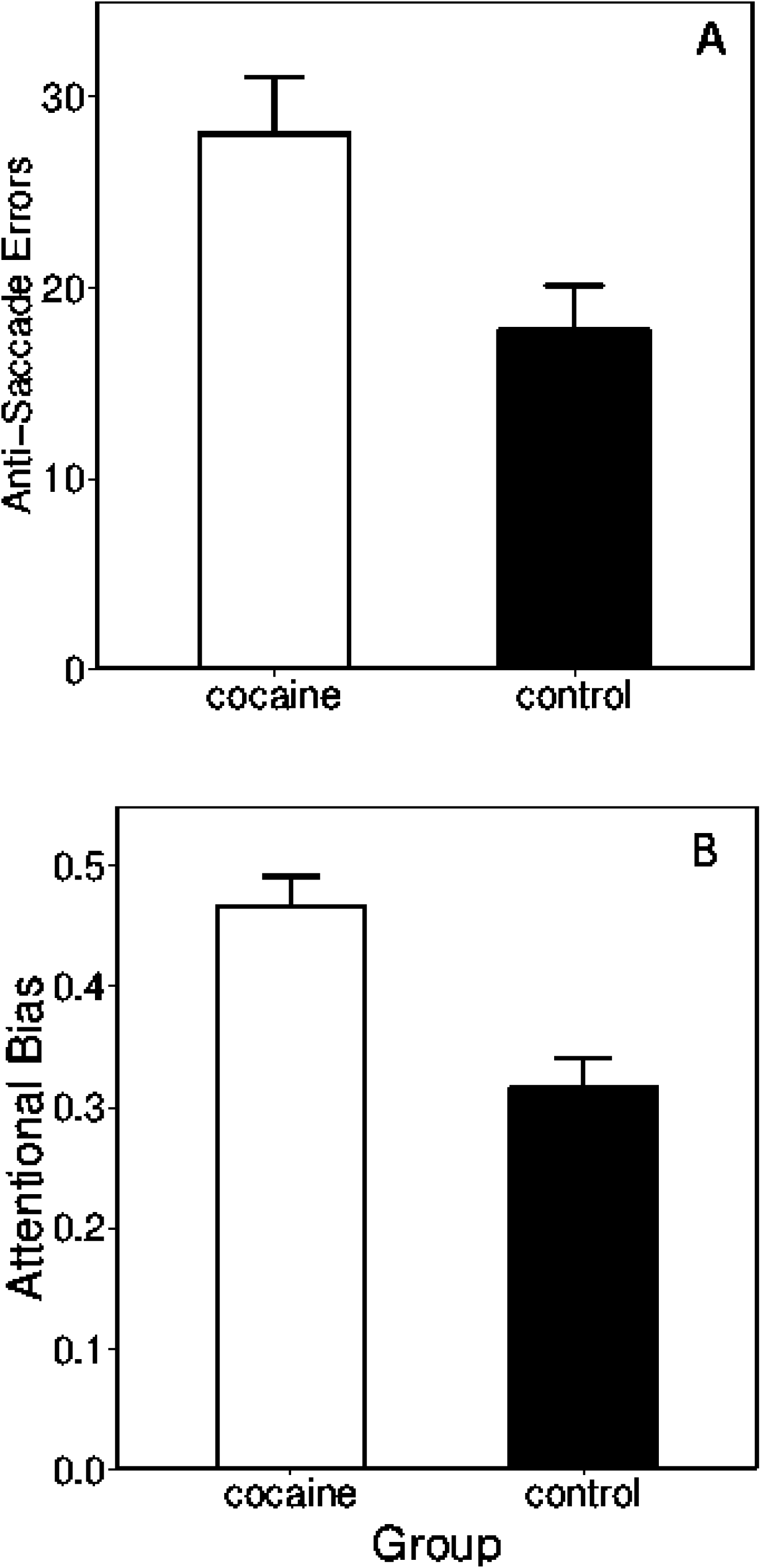
Anti-saccade performance of CUD/AUD subjects and control subjects on the cocaine eye-tracking task. Panels A and B show mean total anti-saccade errors and attentional bias scores (± SEM), respectively. The attentional bias score was calculated as anti-saccade errors on cocaine-stimulus trials / total anti-saccade errors on all trials, providing an indicator of attentional bias towards cocaine cues.

### Anti-Saccade Task: HDDM

The posterior probabilities for the HDDM drift rates (*v*) for the CUD/AUD and control groups are shown in Fig 3, depicting both consistently longer and more highly variable drift rates in CUD/AUD across all stimulus types. Between groups, the Bayesian posterior probabilities that *v* for CUD/AUD > control on cocaine-, neutral-, and shape-stimulus trials were 0.99, 0.94, and 0.95, respectively. Examining *v* among stimulus types within the CUD/AUD group, the following Bayesian posterior probabilities were observed: cocaine > neutral = 0.90; cocaine > shape = 0.96; neutral > shape = 0.68. For the control group, the corresponding probabilities were 0.47, 0.74, and 0.77, respectively. Shown graphically in Fig 3, note that a posterior probability = 0.50 is essentially chance (note overlapping distributions in the control group corresponding to p = 0.47). For the frequentist linear model of drift rate (*v*) *difference scores* (cocaine – neutral+shape stimuli), df = 3, 55, R^2^ = 0.10: group t = 2.08, p < .05. CUD/AUD had higher drift rates than controls, 0.17 (± 0.05) vs. 0.05 (± 0.04).

**Fig 3.**
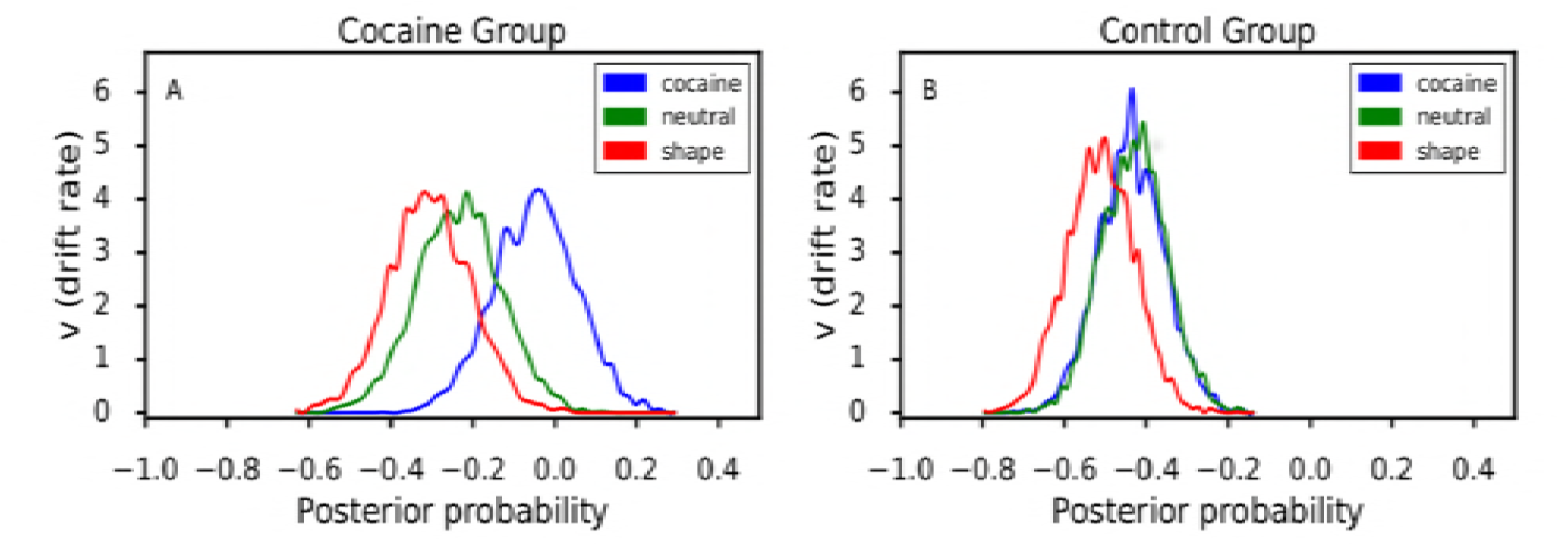
Posterior probabilities of drift rates (*v* parameter) derived from the Bayesian hierarchical drift diffusion model (HDDM) for each stimulus type presented on the anti-saccade task. The HDDM models decision making under two-option conditions; the options here were executing a saccade toward the stimulus or an anti-saccade away from the stimulus. HDDM uses Markov chain Monte Carlo to estimate posterior distributions for all model parameters based on both response accuracy and time to execute the response (e.g., execute a saccade), the results of which are plotted for *v* in Figure 3 (see 59 for HDDM computational details). Drift rate can be conceptualized as the rate at which information is accumulated toward a decision-threshold prior to executing a response. Stimulus contexts presenting, for example, greater conflict or increased background noise should increase drift rate. In Figure 3, drift rate parameters are shown separately for the cocaine group (panel A, left) and the control group (panel B, right).

## Telomere Length

Telomere length was measured as the T/S ratio in a linear model with df = 3, 45, multiple R^2^ = 0.04. T/S ratio of the CUD/AUD group was less than the control group (0.93 (± 0.07) vs. 1.13 (± 0.14), i.e., shorter telomeres) but these differences were not statistically significant (t = −1.16, *ns*).

### Associations with White Matter

As described above, relationships among RD and cocaine and alcohol use, age, anti-saccades, and telomere length were examined with penalized regression using the elastic net with leave one out cross-validation, which established the optimal α and λ parameters based on RMSE. The best fitting model had the following features: α = 0.10, λ = .019, RMSE = 0.13, and R^2^ = 0.79. Utilizing the R selectiveInference package as per [62,63], age, anti-saccades (HDDM difference score), telomere length, and years of cocaine use were all important predictors (*p* < .05), while years of alcohol use was not. Fig 4 depicts the multivariate relationships between RD and the predictors, and Table 2 provides penalized regression coefficients with 95% CI, Z-scores, and *p*-values for each predictor. Table 2 reveals that years of cocaine use and age were the strongest predictors of white matter integrity as indexed by RD. Supplement S4 Data and Figures provide comprehensive graphical and numerical indices of the elastic net regression parameters and model fits.

**Fig 4.**
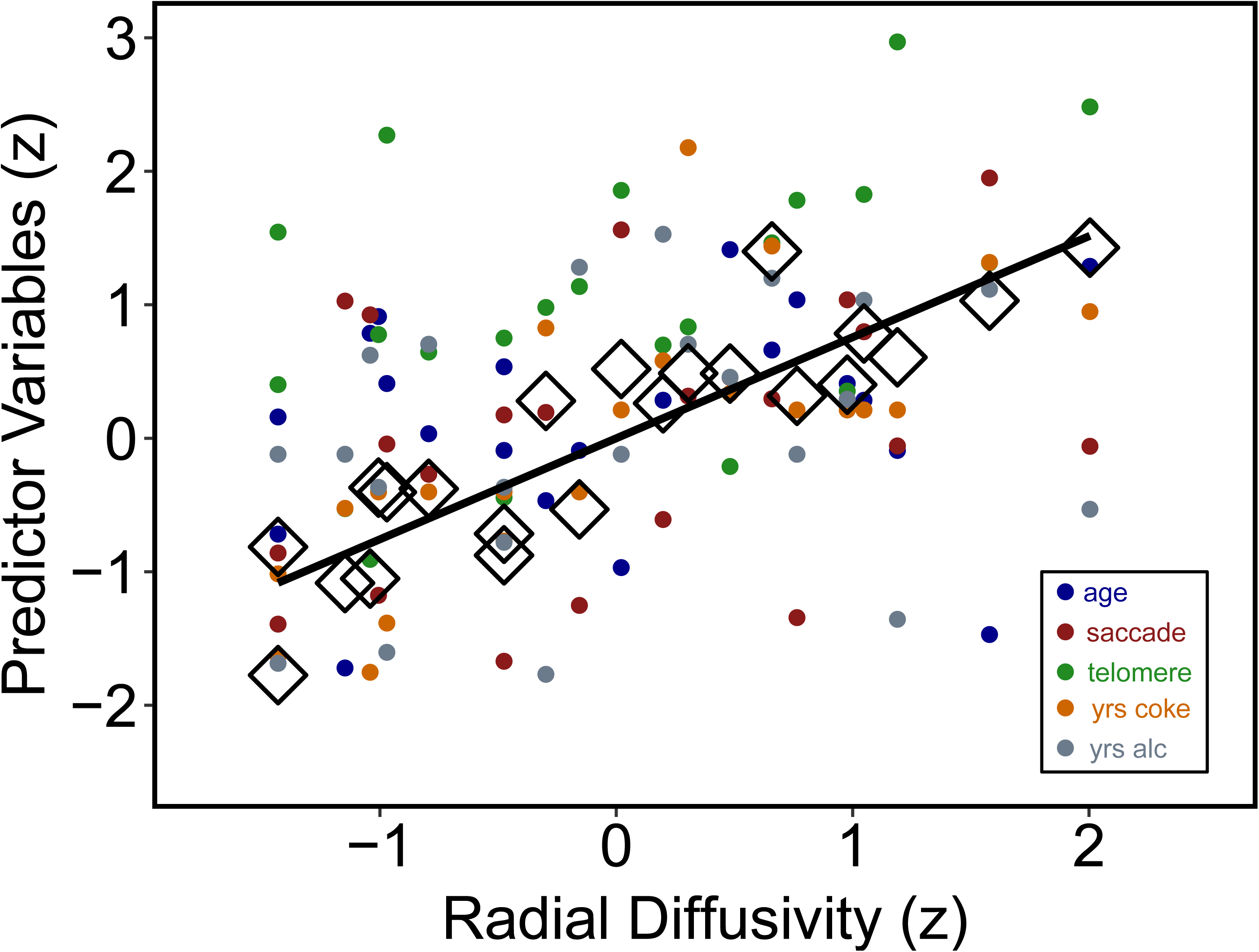
Scatterplot of relationships between radial diffusivity and individual predictors in the elastic net regression model. Within the CUD/AUD group, Figure 4 shows a scatterplot of the relationships between radial diffusivity (x axis) and individual predictors in the elastic net regression model (y-axis): age, anti-saccade drift rate differences (cocaine – non-drug stimuli, see Figure 3), telomere length (T/S ratio), years of cocaine use, and years of alcohol use. Predictors are standardized (z-scored) for graphical interpretability and presentation on a common scale. Large open diamonds (◊) show the predicted points derived from the elastic net model, and the solid black line shows the model-derived line of best fit, R^2^ = 0.79.

**Table 2.** Penalized regression coefficients with 95% CI, Z-scores, and *p*-values for each predictor in the elastic-net regression with DTI radial diffusivity (RD) as the outcome variable.

| Variable | Coefficient (95% CI) | Z-score | $p$-value |
| --- | --- | --- | --- |
| **Age** | 0.375 (0.148 – 0.595) | 2.862 | 0.004 |
| **Anti-saccade:**<br>**Drift rate difference** | 0.327 (0.088 – 0.547) | 2.453 | 0.014 |
| **Telomere length:**<br>**T/S ratio** | -0.330 (-0.560 – -0.004) | -2.363 | 0.047 |
| **Cocaine use (years)** | 0.606 (0.352 – 1.048) | 3.884 | < 0.001 |
| **Alcohol use (years)** | 0.075 (0.592 – -0.721) | 0.540 | *ns* |

## Discussion

In this dataset we showed that chronic cocaine and alcohol use were associated with: (1) WM integrity related to duration of cocaine use in extensive WM tracts, and (2) decreased saccadic control in the presence of cocaine-cues compared to control subjects. Additionally, we found that radial diffusivity was predicted by years of cocaine use, age, anti-saccade performance, and telomere length. While each of the predictors will require verification via systematic replication, one interpretation of the collective results is a (partial) depiction of the allostatic load imparted by chronic cocaine and alcohol use.

Studies of this nature are fundamentally constrained by measurement of a limited set of many potential key variables and the ability to address alternative hypotheses, imposing the requisite conclusion that the findings should be considered preliminary. In addition, limitations in sample size and study design likely prevented uncovering pairwise associations among all these individual predictors, if they are indeed related. Accordingly, one might cautiously presuppose that the present results are related to chronic cocaine use plus the additional burden of alcohol abuse (although alcohol use provided no independent predictive utility), plus the influence of unmeasured factors. Such factors are likely to include variables known to alter neurobehavioral trajectories, e.g., trauma exposure, traumatic brain injury, genotypic variation, and prenatal exposure to abused substances and environmental toxicants [12,16,65-68].

With regard to measurement limitations, markers of HPA-axis, hormone, and immune system activation, as well as hedonic reward deficits are well-established indicators of allostatic load [3,69]. The addition of these markers in the present study would have allowed for analyses of co-variation among established markers and those reported here. Clearly, as work in this area matures, connecting markers like FA/RD and cell aging with markers of HPA-axis, immune system response, and hedonic deficits will advance allostatic load hypotheses in human substance use disorders. One challenge to this aim will be overcoming measurement incongruity. DTI and cell aging (e.g., telomere length) markers tend to be static over the time durations of most experimental work, while HPA-axis and immune responses are dynamic systems subject to phasic changes over short time periods. Nevertheless, bridging these measurement domain stands to advance understanding of allostatic processes in SUD.

Collectively, the data suggest that these variables are possible indicators of CUD/AUD dysregulation in important biobehavioral systems. For chronic SUD, this is consistent with models of allostatis, which “…involves the whole brain and body instead of simply local feedbacks… When demands become chronic, the brain-body system tonically adapts at essentially all levels of organization…” [70]. Notably, duration of cocaine use, anti-saccade performance, and telomere length were all predictors of decreased white matter integrity. Given the study limitations, including the restricted sample, none of the outcomes should be individually considered as strong evidence of allostatic load. However, together the aggregate results suggest a possible allostatic shift associated with chronic cocaine and alcohol use.

Duration of cocaine use and telomere length can be considered indicators of chronicity. Previous studies have shown that severity and/or extent of cocaine use (with polysubstance use) was related to decreased WM integrity [12,71], while abstinence from cocaine was related to specific fiber tract improvements in FA value [35,72]. Associations between CUD and reductions in white matter integrity suggest allostatic shifts and possible neurotoxicity to white matter neurons [11,73,74]. These arguments are not fully verifiable based on the present dataset, but provide intriguing hypotheses for systematic replications in broader samples of participants with CUD/AUD. Because the regression analysis was conducted in an exploratory manner, it should be considered hypothesis generating rather than confirmatory of any relationships specified *a priori*. Moreover, alterations in common WM regions have been observed between substance abuse and other addictions such as gambling [75], raising the alternative explanation of preexisting conditions rather than cocaine-alcohol neurotoxicity. Previous work has established a link between WM impairment (FA, RD) and chronic alcohol abuse [9,13]. The participants in the present dataset had considerably less lifetime alcohol use than typically represented in the alcohol-DTI literature, which may explain why we did not observe independent effects of alcohol use on DTI metrics. Such questions are best addressed by comprehensive longitudinal studies such as the ongoing ABCD project [76].

As AD measures diffusivity running along axons and RD measures the diffusion perpendicular to axons, low AD values have been implicated in internal axonal degeneration [40,41]. Conversely, low RD values have been interpreted as indicators of degeneration of myelin, associated with neurological pathologies, including multiple sclerosis [42], Alzheimer’s [43], and schizophrenia [44]. Indeed, our preclinical work in cocaine-exposed rodents indicates that RD reflects altered myelin integrity [11]. In this context, we observed no relationships between CUD and AD values but significant associations between CUD and RD/FA(R^2^ = 0.51 and 0.56), providing additional support linking CUD to myelin impairment [11, 77]. Whether these findings are related to direct neurotoxic properties of cocaine on white matter fibers, or water-like edema, or interactions with pro-inflammatory mechanisms (glial cell activation, [78]), intra-axonal injury, CNS stress mechanisms (HPA-axis activation, [6]), or dysregulation of myelin-related gene expression (Myelin Basic Protein, [79]) are unknown. In vitro work at cellular and molecular targets may help reveal the role of these putative mechanisms [84].

While T/S ratio was shorter in CUD/SUD participants, we did not observe a statistically significant difference from controls. This may represent a limitation of sample size; studies of telomere length in psychiatric or SUD samples have typically examined larger samples [17,18,26,27,80]. Additionally, while the T/S index is well established, more recent measurement techniques involving DNA methylation and mitochondrial DNA offer potentially greater measurement sensitivity [81,82]. However, within the CUD/AUD group, T/S ratio was negatively associated with RD. Reductions in telomere length may indicate common processes in neurodegenerative disease [83].

Collectively, the results add to a growing literature suggesting that chronic cocaine use – and the risk factors associated with a CUD lifestyle – impart an allostatic load measurable across multiple domains of inquiry. While not part of the stock indices of allostatic load (e.g., HPA-axis dysregulation, hypertension, cell aging, oxidative stress), important domains to be considered for future work include white matter integrity, epigenetic changes, neuroimmune mechanisms, and cognitive integrity.

## Acknowledgements

This work was supported by National Institute on Drug Abuse (NIDA) grants DA P50 009262 (JMS, FGM, SDL, PAN), U54DA038999 (FGM) and UL1TR002649 (FGM), the UTHealth McGovern Scholars Award (SDL), and funds from the Center for Neurobehavioral Research on Addictions at the UTHealth McGovern Medical School (JMS, SDL). The authors wish to thank Vipul Kumar Patel, Jessica Vincent, Edward Zuninga, Joseph Alcorn 3^rd^, and Zahra N. Kamdar for their excellent technical support.

supporting information

